# Taste Bud Connectome: Implications for Taste Information Processing

**DOI:** 10.1101/2021.04.20.440689

**Authors:** Courtney E. Wilson, Robert S. Lasher, Ruibiao Yang, Yannick Dzowo, John C. Kinnamon, Thomas E. Finger

## Abstract

Taste buds contain 3 types of morphologically identifiable mature cells, 2 of which mediate transduction of specific taste qualities: Type III cells transduce sour while Type II cells transduce either sweet, bitter or umami. A long-standing controversy is whether the nerve fibers innervating these cells are wired specifically, in a labeled-line fashion, or non-specifically, leading to broad responsiveness across taste qualities, the so-called across-fiber system of encoding. Using serial blockface scanning electron microscopy through 5 circumvallate mouse taste buds, we reconstructed the patterns of connectivity of nerve fibers as well as the degree of potential interaction between the two types of taste transducing cells. Type II and Type III cells share few points of contact with one another, and display no morphologically identifiable synapses, suggesting limited direct interaction between these cell types. Of the 127 nerve fibers that make synaptic contacts with taste cells in the sampling volume, about 70% (n=91) synapse with only one taste cell. Of the remaining 36 fibers, 4 synapse with both Type II and Type III cells, the remainder synapsing exclusively with multiple Type II taste cells or multiple Type III cells. Since Type II and Type III cells transduce different taste qualities, the few mixed fibers do not follow a labeled-line organization according to taste quality information and show that connectional specificity in taste buds is not absolute.

**Significance Statement:** Taste buds, the sensory end organs for the sense of taste, contain multiple types of sensory cells, with each responding to one of the primary tastes: salt, sweet, sour, bitter and umami. A long-standing question is whether each type of taste cell is wired specifically to a unique set of nerve fibers conveying a “labeled-line” message to the brain. Using serial sections, we determined the neural connectivity in mouse circumvallate taste buds. The majority of individual nerve fibers connect to a single type of taste cell, but 3.1% of the fibers branch to receive input from taste cells known to have different specificities. Thus taste cannot entirely be carried along nerve fibers dedicated to single taste qualities.

## INTRODUCTION

The sense of taste can distinguish at least 5 principal sensory qualities: salt, sweet, sour, bitter and umami (Ohla et al., 2019). The initial detection of these diverse qualities is accomplished by the specialized epithelial cells of taste buds, called taste cells (Roper and Chaudhari, 2017a). Each transducing taste cell responds principally to one of the five taste qualities according to the molecular nature of the receptor employed. The manner in which this information is encoded and transmitted by the sensory nerves remains open to question.

Although the taste cells themselves generally express molecular receptors for only one taste quality (Chandrashekar et al., 2006), the responses of taste cells are not always restricted to a single quality (Caicedo et al., 2002; Chandrashekar et al., 2006; Dutta Banik et al., 2020). Similarly, responses of individual gustatory nerve fibers range from specific, responding to a single taste quality, to multimodal, responding to multiple qualities (Wu et al., 2015). The origin of this broader responsiveness is unclear and may reside in lack of specificity (Dutta Banik et al., 2020) in taste transducing cells, cross-talk between these cells (Tomchik et al., 2007; Roper and Chaudhari, 2017b) within a taste bud, lack of specificity in synaptic connectivity, or all of these.

The “labeled-line” hypothesis of information encoding holds that each particular functional class of taste transducing cell, e.g. sweet, synapses only with a particular subset of quality-specific nerve fibers (Chandrashekar et al., 2006). In this scheme, analysis of the connectivity within a taste bud should reveal specificity, i.e. a particular nerve fiber should receive synapses only from one particular functional type of taste transducing cell. Conversely, the “across-fiber” hypothesis of taste coding states that some nerve fibers may be broadly responsive and that taste quality can be extracted only by comparing activity across fiber populations. This model does not require precise connectivity within a taste bud and allows for substantial cross-talk between taste cells of differing specificities. In this report, we reconstruct the patterns of taste cell-to-nerve connectivity, the so-called “connectome” by means of serial blockface scanning electron microscopy (sbfSEM). The high resolution of this process enables us to identify all sites of synaptic interaction within a taste bud, including taste cell-to-nerve fiber synapses as well as contacts between taste cells themselves indicative of potential information exchange between cells within a taste bud.

Taste cells responsive to sour, Type III cells (Huang et al., 2008; Kataoka et al., 2008; Chang et al., 2010; Teng et al., 2019; Dutta Banik et al., 2020), have molecular and ultrastructural features distinctive from Type II cells -- those responsive to sweet, bitter or umami (Clapp et al., 2004; Chandrashekar et al., 2006; Roper and Chaudhari, 2017a; Yang et al., 2020). Furthermore, these two different types of taste transducing cells have different relationships to the afferent nerves. Type III cells (sour) utilize conventional synapses where a cluster of synaptic vesicles lies apposed to the presynaptic membrane which abuts the postsynaptic nerve process (Kinnamon et al., 1985; Royer and Kinnamon, 1991; Yang et al., 2000). In contrast, Type II taste cells rely on an unconventional synaptic structure (Romanov et al., 2018) characterized by a large “atypical” mitochondrion (Royer and Kinnamon, 1988) closely associated with CALHM1/3 channels (Ma et al., 2018; Kashio et al., 2019) which release the neurotransmitter ATP upon activation of the taste transducing cell (Murata et al., 2010; Taruno et al., 2013). Using these different ultrastructural features as markers of synaptic loci, we were able to map the patterns of connectivity of taste cells to nerve fibers to ask the question of whether any nerve fibers synapse with both Type II and Type III taste cells thereby testing specificity of connectivity within the bud.

## METHODS

Serial Block-face Scanning Electron Microscopy (sbfSEM) [see (Yang et al., 2020)] was used to generate two datasets from adult (>45 days) mouse circumvallate taste buds: DS2, composed of 563 sections, and TF21, composed of 633 sections. The data used in the present study include 6 different taste buds -- three partial taste buds (two of which were conjoined) from DS2, and two complete and another nearly complete taste bud from TF21 (Fig.1). Four of the taste buds of the connectome dataset were included in the data described in (Yang et al., 2020); an additional taste bud described in that work (TF21_TB0) was examined for cell-cell contacts but was excluded from the connectome analysis because it included too few cells. A different taste bud, (TF21_TB2) is now included in the present work but not described in the previous paper. Cells and fibers that were only partially contained in the sample volume and which showed no evidence of synapses were not included in the connectome data, although such cells were cataloged in terms of cell-cell contacts. The image data include a total of 53 Type II cells, 44 Type III cells and 1 indeterminate cell herein termed Type I/II. Only 42 Type II, 33 Type III cells and the Type I/II cell showed synaptic foci within the imaging volume and were included in the connectome. The connectome data include a total of 127 nerve fiber profiles that form synapses with the various taste cells; 61 could be traced fully to a point of exit from the base of the taste bud. The original image data are freely available at the Electron Microscopy Public Image Archive, part of European Bioinformatics Institute (EMBL-EBI), in dataset <u>EMPIAR-10331</u>.

**Fig. 1:**
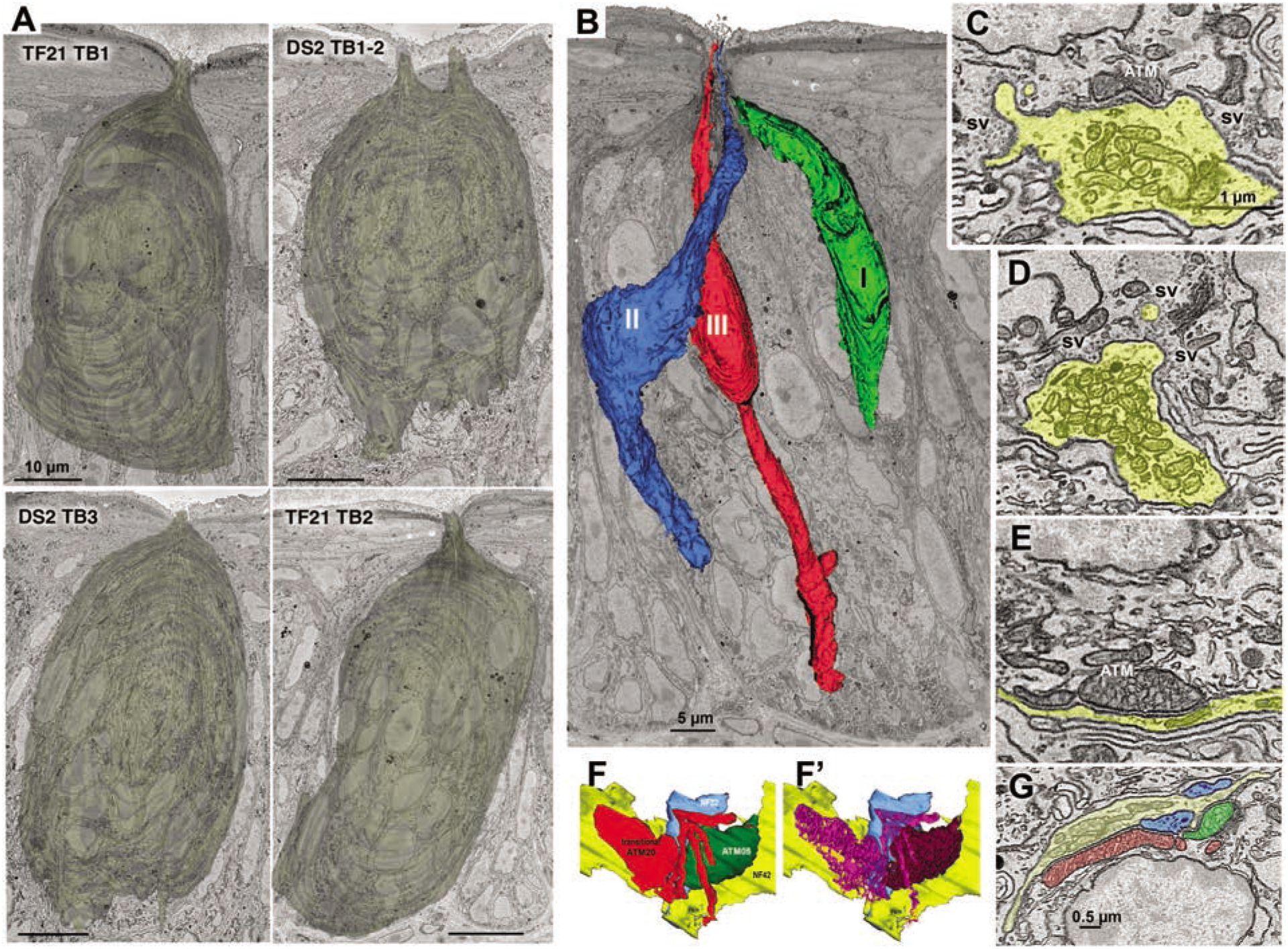
Taste buds, cell types and synapses. **A.** Reconstructions of the appearance of the 5 segmented taste buds in the connectome dataset. The upper right image (DS2_TB1-2) shows the conjoined taste buds with 2 distinct apical pores. Scale bars 10μm. **B**. Taste bud TF21_TB2 showing representatives of the three elongate cell types. **C**. Hybrid type synapse showing both an atypical mitochondrion (ATM) and a cluster of synaptic vesicles (sv) at the region of contact between the nerve fiber (yellow) and Type III taste cell. **D.** Vesicular synapse characteristic of Type III taste cells. Numerous, densely packed conventional mitochondria are containined within the nerve process (yellow). **E.** Channel synapse characteristic of Type II cells with ATM apposed to the plasma membrane at the region of contact with the nerve fiber. **F & F’:** 3-D reconstructions of a transitional mitochondrion at a synapse between a Type II cell and two nerve fibers (blue and yellow). **F** and **F’.** The transitional mitochondrion (red) with mixed lamellar and tubular cristae (magenta, F’) is adjacent to an ATM (green) containing only tubular cristae (dark red, F’). **G.** EM image of the mitochondrial complex reconstructed in panel F.

The two conjoined small taste buds of DS2 (containing within the connectome map 5 Type II cells and 4 Type III cells for TB1; and 4 Type II cells and 2 Type III cells for TB2) were fused along the basal lamina, although each taste cell extended an apical process into only one or the other taste pore. Cells were allocated to these conjoined buds according to which taste pore the cell was associated with. Nerve fibers innervating the taste cells of these conjoined taste buds generally innervated cells in one or the other taste bud, but two nerve fibers received synapses from Type III cells associated with the two different buds.

Every taste cell in each section was assigned a unique identifier and was analyzed using Reconstruct software (Synapse Web Reconstruct, RRID:SCR_002716) (Fiala, 2005) to determine cell type, and whether Type II and Type III cells were in contact. When such contacts were identified, the segmented measuring tool in image J (National Center for Microscopy and Imaging Research: ImageJ Mosaic Plug-ins, RRID:SCR_001935) (Schneider et al., 2012) was utilized to measure the length of each contact and ultimately determine the surface area (using the measured contact lengths and the overall number of sections showing the contact). For taste cells with contacts spanning fewer than 10 sections, the length of each contact in every section was measured. For taste cells with contacts spanning more than 10 sections, the contact length was measured in every fifth section to calculate the surface area of the contact.

The defining characteristics of synapses from Type II cells in mouse circumvallate taste buds are the presence of large, “atypical” mitochondria (ATMs), with tubular cristae, closely apposed to the plasma membrane of the Type II cells at the area of contact with a nerve fiber (Fig. 1E, G) (Romanov et al., 2018; Yang et al., 2020). The plasma membranes of both the Type II cell and nerve fiber are slightly thickened and maintain a constant spacing between them. Similarly, the external membrane of the ATM maintains a fixed distance from the plasma membrane of the taste cell often with patches of dense material bridging this gap.

As viewed with transmission electron microscopy, synapses associated with Type III cells in mouse circumvallate taste buds e.g., (Kinnamon et al., 1985; Yee et al., 2001; Yang et al., 2020) have many of the characteristics of synapses seen in the mammalian central nervous system (Peters et al., 1991; Harris and Weinberg, 2012) including clusters of 40-50 nm vesicles in the presynaptic partner at the point of contact with the postsynaptic fiber, and a slight increase in the density of the pre- and postsynaptic membranes associated with the synaptic cleft (Fig. 1D). However, the increase in the density of pre- and postsynaptic membranes is often not evident using sbfSEM. In addition, at the low voltages used (~2.2 KV) for scanning, only about the top 30 nm of the 75-80 nm thick section is imaged, so synaptic vesicles deeper in the tissue section may not be visible or recognizable. Identification of other types of junctions such as gap junctions would also be problematic because of this limitation on resolution and specimen preparation techniques.

In both datasets, we find a type of specialized, non-synaptic contact between nerve fibers and taste cells which has not been described previously. Based upon its appearance, we term this a “wavy junction”. This type of contact is unlike a gap, macula adherens or desmosomal type of junction. In wavy junctions, closely associated plasma membranes of taste cells and adjacent nerve fibers separate to form a gap of approximately 40-50 nm in the region of apposition. The plasma membranes have the appearance of waves in the regions of contact (arrows, see Figure 2). The contacts can be extensive, sometimes several microns in length and may extend through approximately 10 sections (750-800 nm). This type of junction can occur between a nerve fiber and each of the various types of taste cells as shown in Fig. 2. We do not consider these to be a form of synapse and therefore have not included these wavy junctions in our connectome map.

**Fig. 2:**
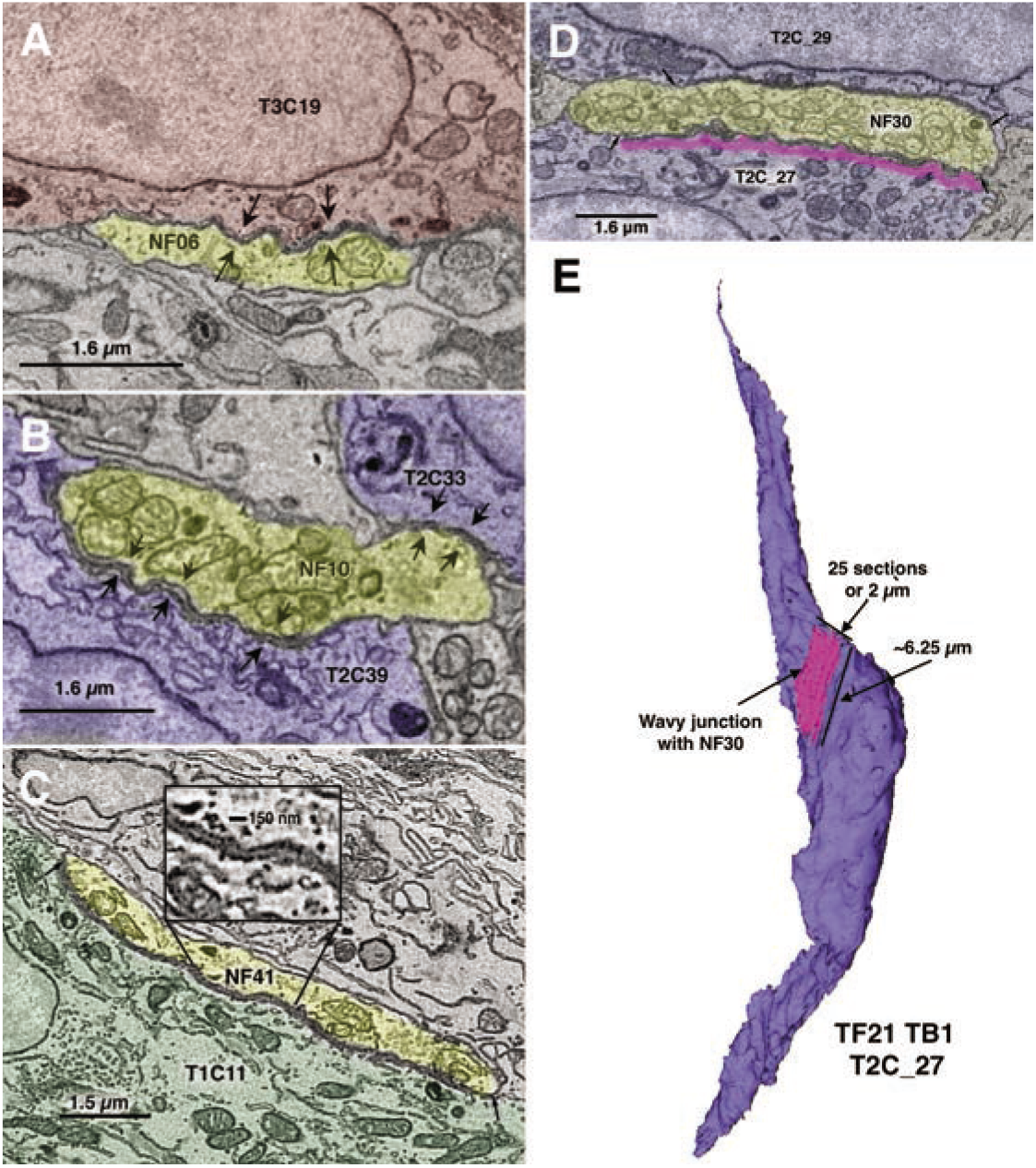
Examples of wavy junctions between nerve fibers (yellow) and diverse cells of taste buds. **A**. From TF21_TB2 : Wavy junction between a nerve fiber (NF06, yellow) and a Type III cell (T3C19, red). Arrows mark the extent of the contact, in this case about 3 microns. **B.** From TF21_TB1: Wavy junction between a nerve fiber (NF10, yellow) and two Type II cells, T2C39 and T2C33, blue. Arrows mark the extent of the contacts, in this case about 1 micron for T2C33 and about 3.5 microns for T2C39. **C.** From DS2_TB3: Wavy Junction between a nerve fiber (NF41, yellow) and a Type I cell (T1C11, green). The inset shows an enlargement of the gap region of approximately 40-50 nm. Arrows mark the extent of the contact, in this case about 8 microns. D. A large wavy junction from TF21_TB1 cell T2C27. Magenta shows the extent of the wavy junction with NF30. E. Reconstruction of cell T2C27 and its wavy junction with NF30 corresponding to the single plane image in D.

Cells and nerve fibers (NFs) were segmented by several individuals; any questionable identifications were resolved by referral to a panel of 2 or more experienced segmenters for consensus. NFs to segment were chosen in several different ways: by random selection from fascicles in the lamina propria, by random selection of nerve profiles from within taste buds, and by tracing back starting from identified synaptic contacts with taste cells. These contacts included both classical vesicle-type synapses and channel synapses (Taruno et al., 2021) marked by an ATM. In addition, NFs partially or completely embedded in cells or closely associated with nuclei were noted and traced. All observations involving interactions between cells and NFs were recorded in spreadsheets for later analysis. All contacts with nerve fibers were checked by at least one other person not doing the original segmentation.

A limitation of the current dataset is that many nerve fibers could not be followed to their origins below the taste bud outside of the sampling volume. The connectome data included in this report identify all nerve fibers that formed apparent synapses with taste cells, but does not include nerve processes which formed no apparent synaptic connections. In three taste buds where we identified all nerve fiber processes, we found approximately half of the nerve fiber profiles formed synapses (DS2_TB1-2: n=67, 53.7%; DS2_TB3: n=61, 45.9%; TF21_TB1: n=62, 43.5%). The nerve processes lacking synapses may have been collaterals of fibers that did form connections with taste cells outside our sampling volume or may represent extending or retracting fibers as identified in other studies (Zaidi et al., 2016).

### Data analysis and construction of connectomes

Recorded observations were collated in a summary sheet and then used to construct preliminary connectomes using mind mapping software (MindNode or iThoughtsX running on a Macintosh computer). Final connectome diagrams were constructed using Adobe Illustrator (RRID:SCR_010279).

### Statistics

Counts of synapses or ATMs in cells were tested using a Chi-square analysis and shown to be not random, i.e. synapses per cell was skewed toward lower values than would occur randomly. The relationship between synapses per cell and number of synapses per fiber was tested using Cohen’s d. Estimation statistics and calculations are taken from estimationstats.com (Ho et al., 2019). For these tests, 5000 reshuffles of the control and test labels were taken; the confidence interval is bias-corrected and accelerated.

## RESULTS

Our analysis of taste bud connectivity included 5 complete or partial taste buds from the circumvallate papillae of 2 mice. The classification of cells as Type I, II and III (Fig. 1) followed classical criteria as summarized in (Yang et al., 2020). In brief, Type I cells have an irregular, multiply-indented nucleus with substantial heterochromatin, and have numerous, characteristic lamellate processes enveloping other cells and nerve fibers. Type I cells terminate apically in a collection of short bushy or arboriform microvilli (Yang et al., 2020). Type II cells have a large, round nucleus with sparse heterochromatin and electron-lucent perinuclear cytoplasm. Apically the cytoplasm is more electron-dense and exhibits substantial distended smooth endoplasmic reticulum. Type II cells included in the connectome display features of a channel synapse, i.e. one or more ATMs apposed to the plasma membrane at the point of contact with a nerve fiber (Romanov et al., 2018; Taruno et al., 2021) (Fig. 1E). Some of these ATMs were continuous with conventional mitochondria with parallel cristae and were interpreted as being transitional (Fig.1F). Nearly all Type II cells terminate apically in a single stout apical microvillus (0.2-0.3 μm width), but we identified 6 of a total of 42 Type II cells, herein termed bushy-ended Type II cells, with multiple microvilli not reaching the taste pore (Fig. 3A). In addition, we found one cell with characteristics of both Type I and Type II cells which we classify as a Type I/II cell (Fig. 3B). This cell has an atypical mitochondrion at the point of apposition with a nerve fiber and an indented nucleus, but lacks lamellate processes typical of a Type I cell. Apically, this cell terminates in a tuft of microvilli indistinguishable from bushy Type I cells described previously (Yang et al., 2020). Since both the bushy-ended Type II cells and the Type I/II cell exhibit an ATM at a point of contact with a nerve fiber (Fig. 3), they are included in the connectome description and map, although we have no evidence that the area of mitochondrial specialization in these cells forms a functional synapse.

**Fig. 3:**
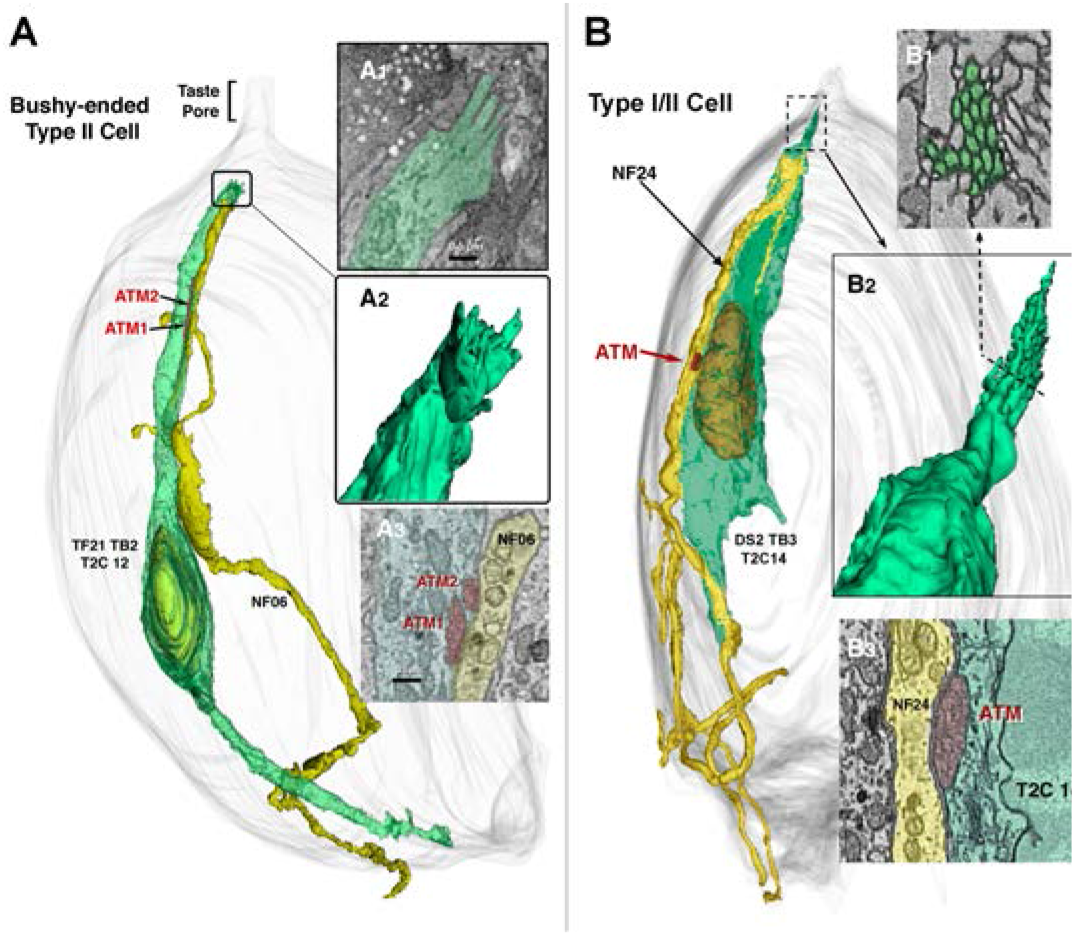
Unconventional Type II cells. **A.** A bushy-ended Type II cell forms an apparent channel type synapse with 2 ATM with nerve fiber NF06 (A3). The apex of this cell (A1-2) ends below the taste pore region in an irregular group of stubby microvilli. **B.** A Type I/II cell shows features typical of both Type I and Type II cells. The nuclear structure and presence of apparent channel synapse with ATM (B3) are indicative of Type II cells but the tuft of apical microvilli (B1-2) are typical of Type I cells.

Type III cells are slender, with a smooth spindle-shape outline, evenly dense cytoplasm with smooth ER lacking swellings, and an elongate nucleus with one or more deep invaginations. Type III cells terminate apically in a single, long microvillus (0.3-0.4 μm width). Type III cells form conventional synapses with nerve fibers including a small collection of presynaptic vesicles (Fig. 1D). A minority (11 of 33) of Type III cells have hybrid synapses with both synaptic vesicles and an atypical mitochondrion (Fig. 1C) as described previously (Yang et al., 2020). These mitochondria with tubular cristae in the Type III cells tend to be smaller than the ATMs of the Type II cells but display the same fixed, close spacing with the plasma membrane as do the ATMs of the Type II cells (c.f. Fig. 1C to 1E & G). The patterns of connectivity of the Type III cells with hybrid synapses do not obviously differ from that of Type III cells with only conventional vesicular synapses but are identified in the connectome maps.

### Cell-Cell Contacts

Although we emphasize relationships between nerve fibers and the known transducing cells of taste buds (Type II and Type III), we also re-examined points of contact between Type I cells and nerve fibers and between Type I cells and the other taste bud cell types. We observed no membrane specializations between Type I cells and nerve fibers, other than wavy junctions (Fig. 2), although as reported previously (Yang et al., 2020), the Type I cells often envelop nerve processes within the taste buds, and nerve fibers often closely approach the nuclei of Type I cells. Multiple examples of intimate, but non-synaptic perinuclear contact between Type I cells and nerve fibers were evident in each taste bud examined.

To test for evidence of cell-cell synaptic communication within the taste bud between Type II and Type III cells that might underlie modulation of taste bud function (Roper, 2013, 2021), we examined areas of direct contact between such heterotypic cells. We also noted rare examples of homotypic contact between taste cells of the same type, i.e. Type II-Type II contacts and Type III-Type III contacts. Nearly all Type II-Type II contacts involved small (<1μm^2^) areas of membrane and exhibited no form of synaptic specialization. A few areas of Type III-Type III cell contact were somewhat more extensive and occasionally involved interdigitation of membrane elements, but exhibited no signs of synaptic transmission such as ATMs, synaptic vesicle clusters or membrane thickenings (Fig. 4).

**Fig 4:**
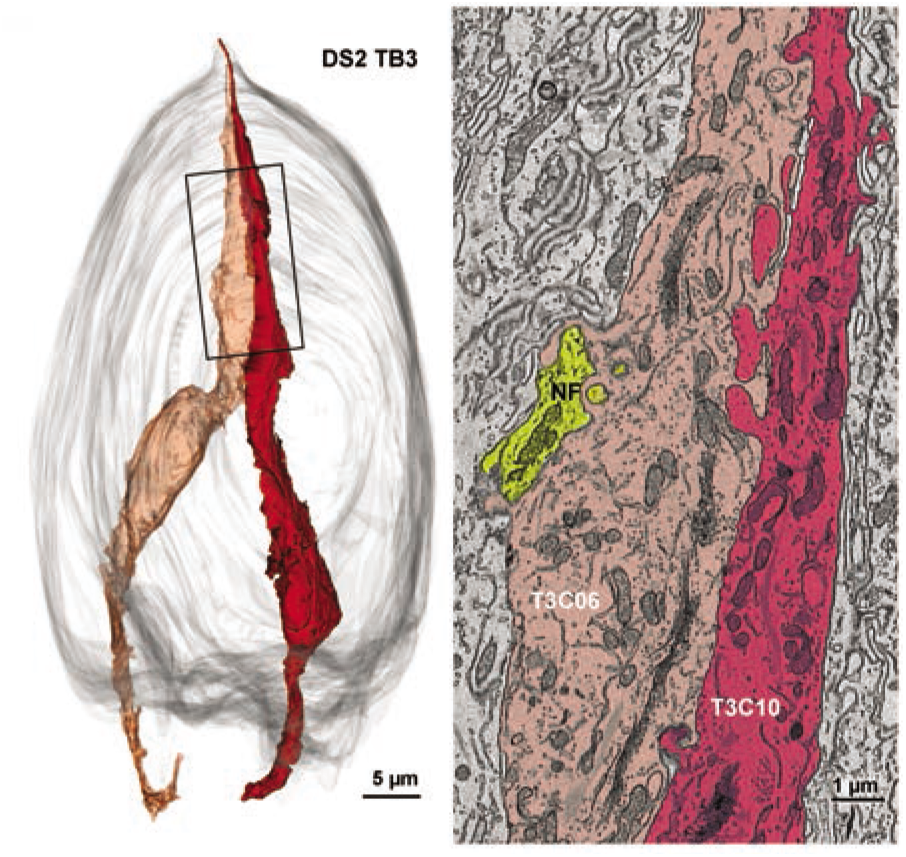
Interdigitating contact between 2 Type III cells. **Left:** 3-D reconstruction of the two Type III cells that form an extensive contact in the apical region of the taste bud. **Right:** Colorized EM image of the region of contact also showing the presence of a nerve fiber (yellow) near the line of contact.

In the 6 taste buds examined in the current study, we find only small, non-specialized points of contact between Type II and Type III taste cells with the areas of contact occurring largely in the upper one-third of the bud and especially in the vicinity of the taste pore where the apices of the different cells converge. Out of a total of 53 Type II cells and 44 Type III cells in the sampling volumes, we find 26 points of contact between Type II and Type III cells involving 16 Type II cells and 17 Type III cells with 4 cell pairs contacting one another at more than 1 point. Of these 26 contact points, 1 lies in the taste pore, 15 lie apically in the upper third of the taste bud, 6 lie centrally and 4 are in the lower third of the taste bud.

These points of contact exhibit no membrane specializations or accumulations of vesicles adjacent to the membranes of either cell. The zones of contact are small, the 2 largest, situated at the apical quarter of each cell, involve about 2 μm^2^ of membrane; 13 of the contacts are less than 0.2 μm^2^ (Fig. 5). The cells exhibit no particular organelles or membrane thickenings along these points of contact. The areas of contact are relatively remote from the synaptic sites from both Type II and Type III cells. These findings suggest that direct cell-cell synaptic communication between Type II and Type III cells does not play a role in taste bud function although non-synaptic interactions, e,g, diffusion of neurotransmitter away from a synaptic site, may be functionally important (Dando and Roper, 2009; Roper, 2013). Technical limitations on the resolution obtainable by this technique do not entirely rule out the possibility that some of these contacts might reflect small, unconventional junctional complexes (Akisaka and Oda, 1978) underlying possible coupling of taste cells (Roper, 2021).

**Fig. 5:**
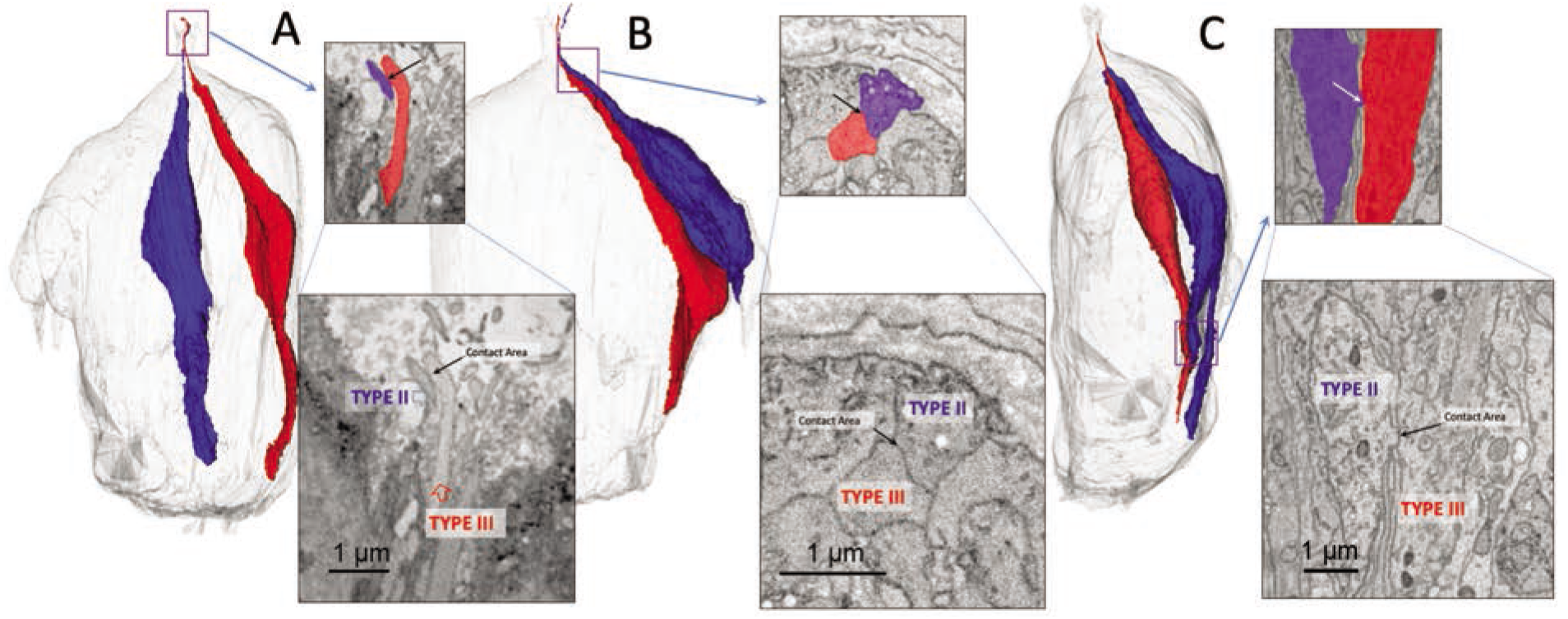
Examples of limited regions of contact between Type II and Type III cells. **A.** Contact between apical processes in the taste pore. **B.** Apical contact below the taste pore. **C.** Contact in the basal region of the taste bud.

#### Taste Cell-Nerve Fiber Contacts

We identified synapses between taste cells and nerve fibers based on the criteria described above, i.e. ATMs closely apposed to the plasma membrane marking channel type synapses (Taruno et al., 2021) for Type II cells and vesicle clusters for conventional vesicular synapses in Type III cells (Fig. 1). The post-synaptic nerve fibers were identified and traced to produce a taste bud connectome for each taste bud. The majority (65.4%) of post-synaptic nerve fibers in the dataset connected to Type II cells (Table 1) while only a few (3.1%) received synapses from both Type II and Type III cells.

**Table 1:** Fiber Innervation of Type II and Type III Cells

|  | Total # | # | % | # | % | # | % |
| --- | --- | --- | --- | --- | --- | --- | --- |
| **TF21_TB1** | 35 | 23 | 65.7% | 11 | 31.4% | 1 | 2.9% |
| **TF21_TB2** | 27 | 17 | 63% | 8 | 30% | 2 | 7.4% |
| **DS2_TB1&2** | 30 | 18 | 60% | 12 | 40% | 0 | 0% |
| **DS2_TB3** | 35 | 26 | 74.3% | 9 | 25.7% | 1 | 2.9% |
| **TOTALS** | **127** | **84** | **66.1%** | **40** | **31.5%** | **4** | **3.1%** |

Conventionally, nerve fibers associated with taste buds are divided into an intragemmal group, within the taste bud, and a perigemmal group which closely surround the taste bud, e.g. (Zaidi et al., 2016). We find occasional fibers which enter the taste bud as intragemmal fibers to receive a synapse from one or more taste cells and then exit the taste bud more superficially to terminate as free nerve endings in the vicinity of the taste pore as reported recently by others for taste buds in fungiform papillae (Huang et al., 2021) (Fig. 6A & B). Conversely, other fibers enter the perigemmal epithelium but continue on to penetrate the taste bud and receive synapses from one or more taste cells (Fig. 6C).

**Fig 6:**
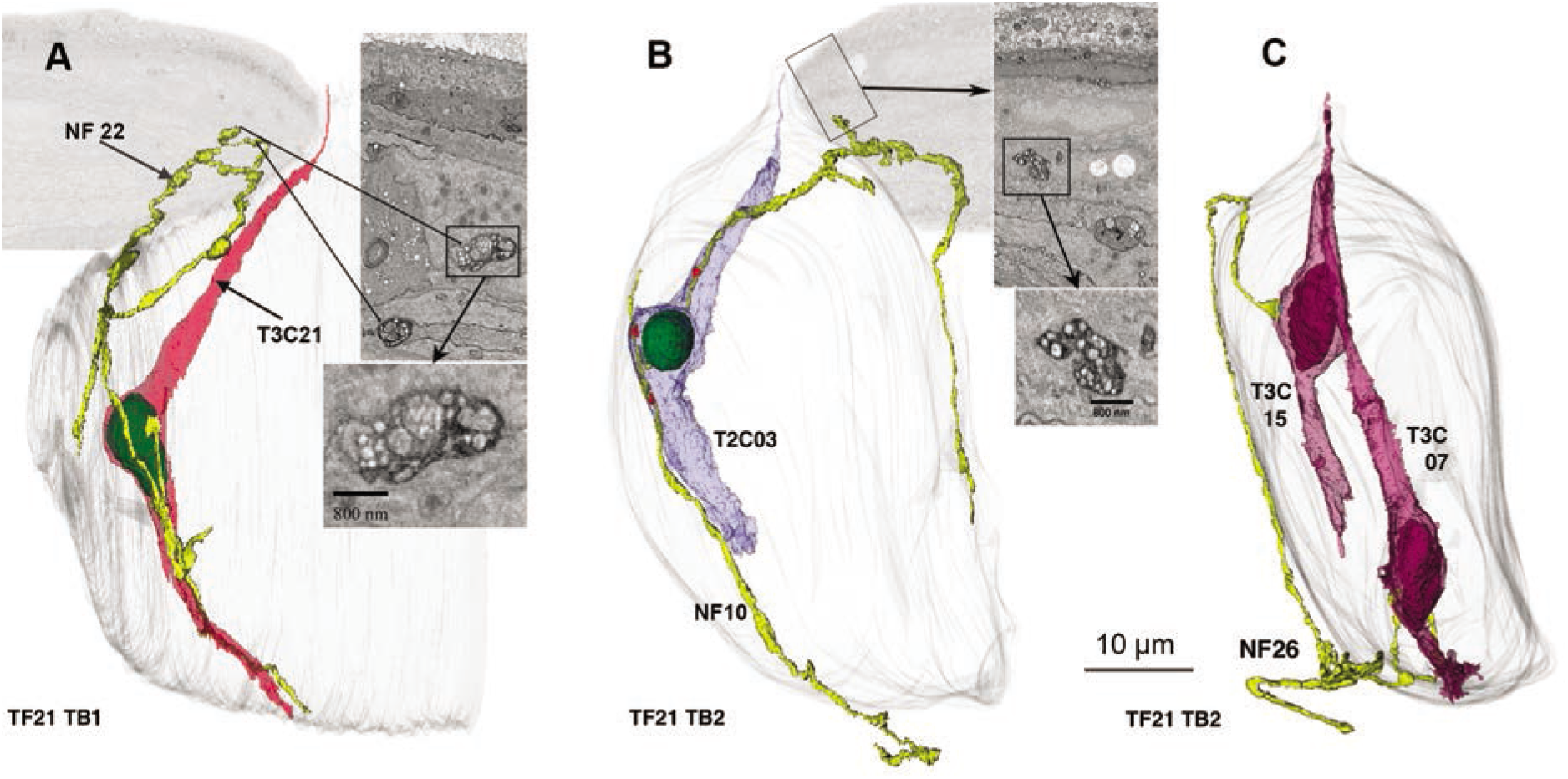
Examples of nerve fibers (yellow) that both synapse with taste cells and traverse perigemmal space. **A.** 3D reconstruction of a fiber that starts as a an intragemmal fiber, synapses with a Type III cell, then exits the taste bud to run in perigemmal space. Inserts at right show EM morphology as indicated. **B.** Similar nerve fiber that synapses with a Type II cell. Adjacent EM images illustrate the nature of the synapses and free endings in the perigemmal epithelium. **C.** An example of a fiber that starts as a perigemmal fiber and then enters the taste bud to form a synapse with 2 Type III cells.

In most, but not all cases, when nerve fibers intimately contact Type II or Type III cells, the taste cells synapse onto the nerve fiber. We do, however, find many examples where a Type II or Type III taste cell contacts a nerve fiber, but no synapse is evident. Such non-synaptic contacts occur without regard to cell type, e.g. a nerve fiber receiving a synapse from a Type II cell may contact other Type II cells without evidence for a synapse, or may contact Type III cells without evidence for a synapse. Fig. 7 shows an example of a nerve fiber receiving a channel synapse from a Type II cell and then extending downward to become enveloped by a Type III cell but without any evident membrane specialization or synaptic vesicles. The absence of a membrane specialization at such points of contact does not, however, preclude exchange of information at these junctions, e.g. by ephaptic interaction. We did not include such points of nonspecialized contact in our connectome map.

**Fig 7.**
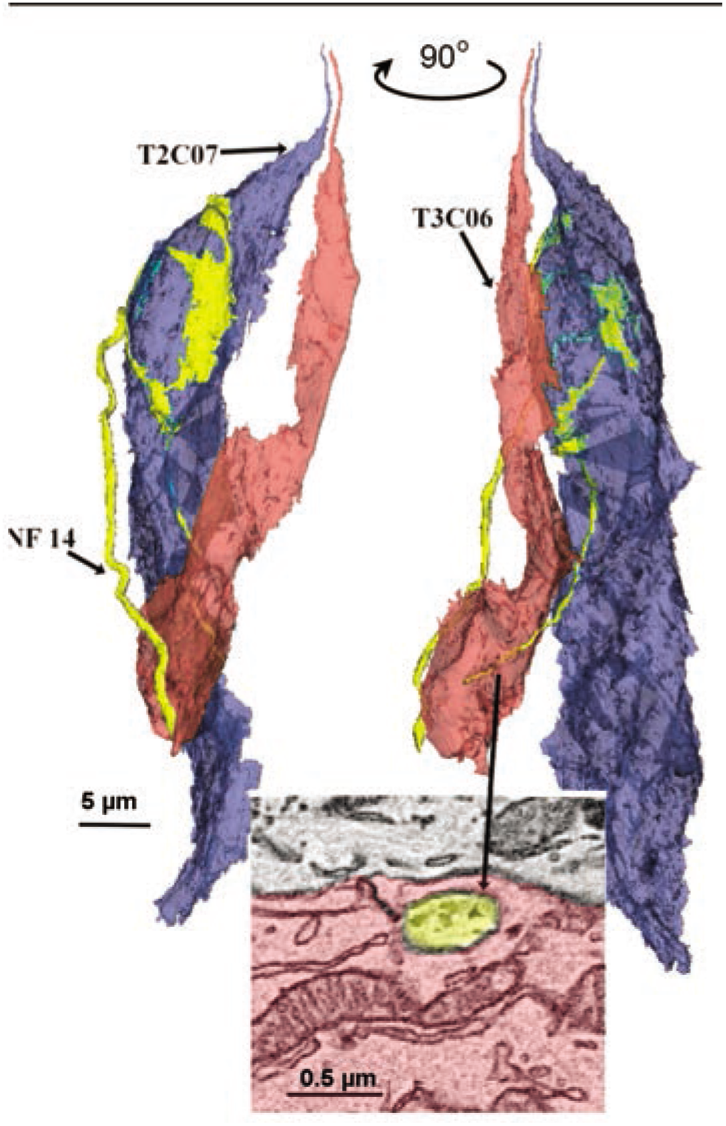
Nerve fibers that contact taste cells do not always form synapses. In this example, showing 2 views rotated about 90°, the nerve fiber (yellow) synapses with a Type II cell (blue) and then runs downward to become fully surrounded by a Type III cell, (see inset below) but without forming a specialized contact with that cell.

A noteworthy feature of taste cell-nerve fiber synapses from both Type II and Type III cells was the variety in number of synapses made by each taste cell and by each nerve fiber (Fig. 8). For Type II cells, the number of channel synapses made by individual taste cells ranged from 1 to over 20, with an average of 6.2 ATMs/cell. The number of synapses per cell for Type III cells (mean = 3.8) was also variable but was significantly lower than the number of synapses per Type II cell. The unpaired Cohen’s d between vesicular synapses (Ves Syn) and channel synapses is 0.52 [95.0%CI 0.102, 0.888]. The p value of the twosided permutation t-test is 0.029.

**Fig. 8.**
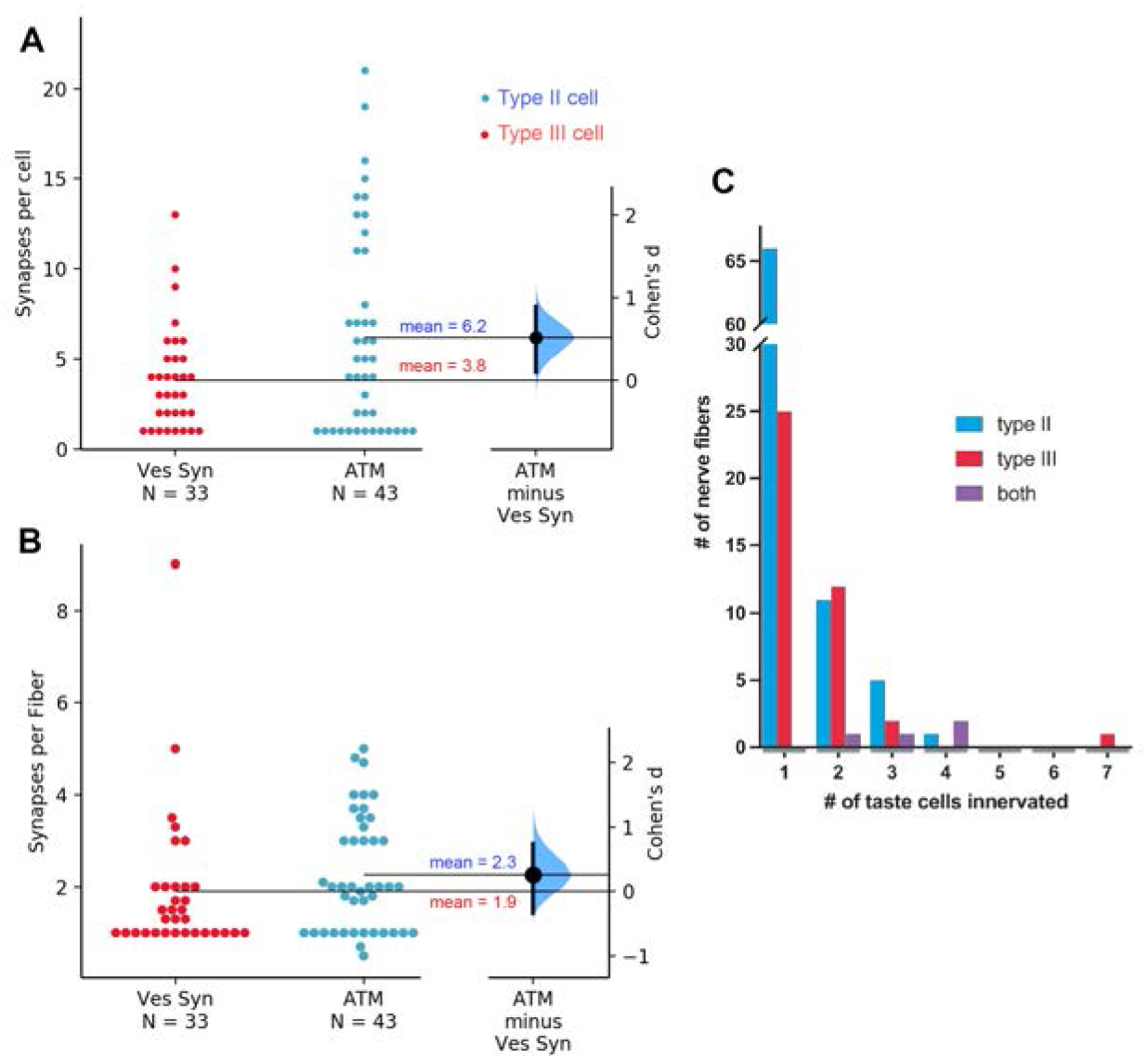
**A**: The mean number of synapses per cell is larger for Type II cells (ATM) than for Type III cells (Ves Syn). The Cohen’s d between Ves Syn and ATM is shown in this Gardner-Altman estimation plot. Both groups are plotted on the left axes; the mean difference is plotted on a floating axes on the right as a bootstrap sampling distribution. The mean difference is depicted as a large dot; the 95% confidence interval is indicated by the ends of the vertical error bar.**B**: Similar plot showing no significant difference between average number of synapses per fiber for Type II (ATM) and Type III (Ves Syn) cell synapses. **C.** Histogram showing how many nerve fibers innervate different numbers of taste cells. Number of nerve fibers innervating Type II cells are in blue, those innervating Type III cells are in red, and those innervating both Type II and III cells are in purple. **C.** Histogram showing how many nerve fibers innervate different numbers of taste cells. Number of nerve fibers innervating Type II cells are in blue, those innervating Type III cells are in red, and those innervating both Type II and III cells are in purple.

Of the 43 fully characterized Type II cells (including the Type I/II cell) in our data, 16 showed only 1 or 2 ATMs) whereas 4 cells had 15 or more. Cells with higher numbers of ATMs tended to be connected to multiple nerve fibers and, even omitting the trivial case for 1 ATM, the number of postsynaptic nerve fibers was proportionate to the number of ATM in any given cell (*r*(30) = .799,*p* <0.001). Often, a single nerve fiber made repeated synaptic contacts onto a single Type II cell, e.g. in TB3 of the DS2 dataset, a nerve fiber (NF04) contacts Type II taste cell (T2C1) at 9 distinct sites. Although Type III cells also displayed multiple synapses with some nerve fibers, the number of such contacts usually was much lower than in Type II cells (1-4 sites per cell). One exceptional Type III cell (T3C2 in TF21_TB1) showed 13 distinct synaptic loci onto 4 different nerve fibers, but the average number of synapses per Type III cell was 3.8 (+/-2.9 s.d.). The number of nerve fibers innervating a particular Type III cell was not significantly correlated with the number of synapses from that cell (r(16) = .30 = p>0.05) although individual nerve fibers often received several synapses from the same Type III cell. Conversely, some cells of both types synapse onto many nerve fibers (see below).

#### Patterns of Connectivity

Within the sampling volume of the largest taste bud in our datasets (TF21_TB2), containing about 100 elongate taste cells (including 12 Type II and 12 Type III synaptically connected cells), we identified 27 independent nerve profiles receiving synapses from taste cells of which 18 fibers could be traced fully until their point of exit from the taste bud. Across all taste buds, the proportion of fibers branching to innervate different taste buds was not significantly different comparing fully traced fibers, i.e. those traced to a point of exit from the base of the taste bud, to truncated nerve fibers, i.e. those traced only until the end of the dataset but still within the taste bud profile (Fibers innervating Type II cells: Full: 8 branched, 28 unbranched; Truncated: 8 branched, 40 unbranched: Chi-square test; p > 0.05. Innervating Type III cells: Full: 7 branched, 17 unbranched; Truncated: 7 branched, 8 unbranched Chi-square test; p > 0.05). An average of 22.7% of fibers innervating Type II cells branch to innervate multiple cells while 41.9% of fibers innervating Type III cells branch (Chi-square test; p > 0.05). As mentioned above the total number of nerve profiles does not necessarily equal the total number of independent nerve fibers entering the taste bud since many of the fibers likely branch below the basal lamina in the subgemmal plexus (Kanazawa and Yoshie, 1996). Nonetheless, we discern several distinct patterns of connectivity as described below.

With a few exceptions, nerve fibers that synapse with a Type III cell synapse only on other Type III cells; fibers that synapse with a Type II cell generally synapse only with other Type II cells (Fig. 9A & 10). Each nerve fiber generally forms synapses with a small number of taste cells – typically one, two or three cells, but some fibers innervate 4 or more cells of the same type (Fig. 9A, B). In the case of Type II cells, 42.9% (18 of 42) are innervated by unique nerve fibers that synapse with no other cells; for Type III cells, 5 of 33 (15.2%) are uniquely innervated. As can be appreciated in the connectome maps (Fig. 10), the remaining cells of each type within a taste bud are interconnected in various ways, especially in the larger taste buds. For example, in TF21_TB2, 9 of the 12 Type II cells synapse onto a network of 7 branched nerve fibers. In DS2_TB3, 5 of 8 Type II cells synapse onto the same pool of 4 nerve fibers. Similarly, Type III cells may synapse onto a connected network, e.g. in DS2_TB1, the 4 Type III cells all synapse onto the same set of nerve fibers; in DS2_TB3, 4 of 5 Type III cells converge onto 3 nerve fibers; and in TF21_TB2, 7 of 12 Type III cells converge onto 2 nerve fibers while 3 other cells converge onto another 2 nerve fibers. Conversely, a single cell may synapse onto numerous fibers (Fig. 9C, D).

**Fig: 9:**
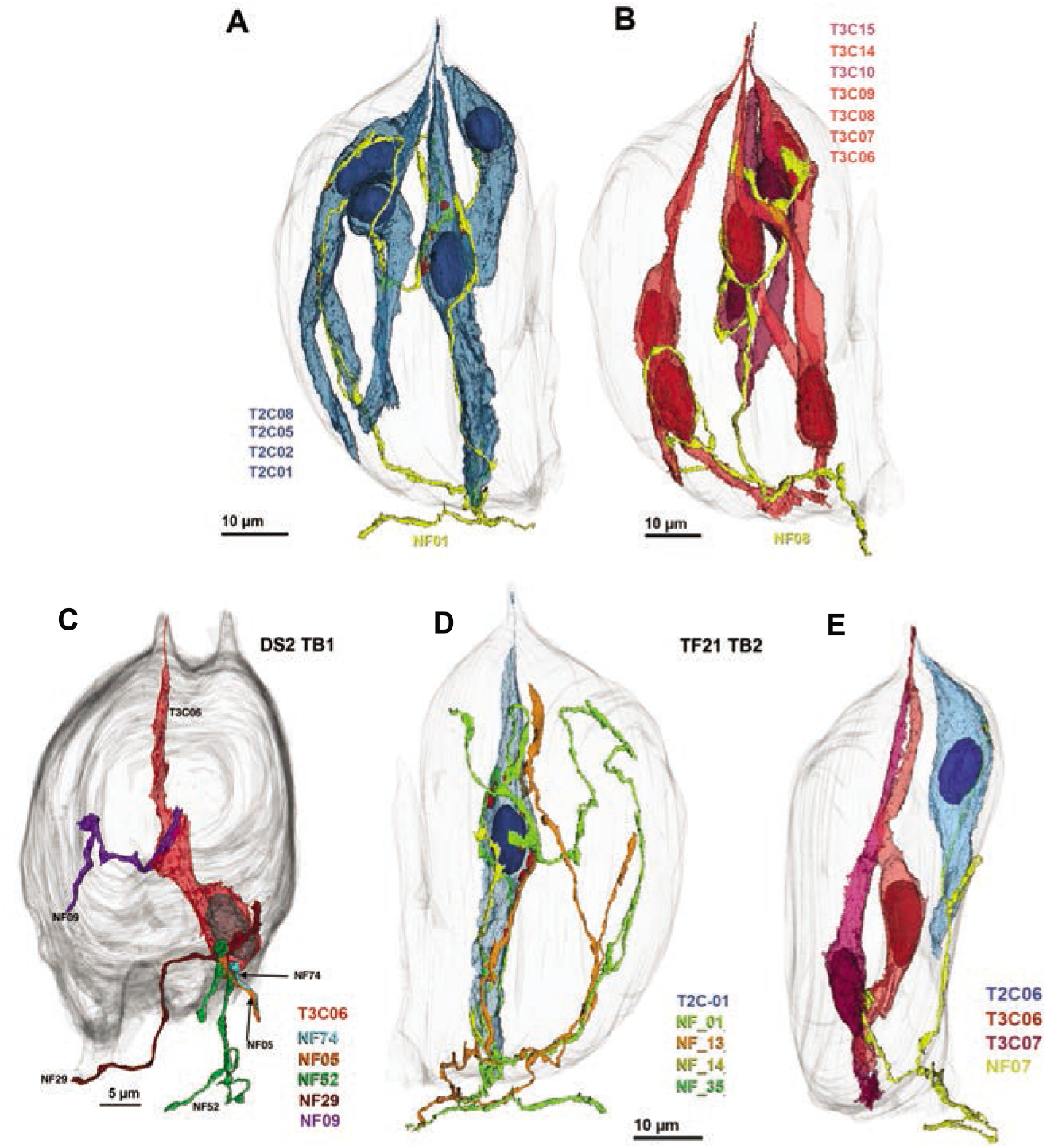
Examples of different patterns of innervation. **A & B.** Individual nerve fibers may connect to many cells of the same type. **A**. Network of 4 Type II cells (blue) connected to a single nerve fiber (yellow). ATM in red. **B.** Network of 7 Type III cells connected to a single nerve fiber. **C & D.** Individual taste cells innervated by multiple nerve fibers. **C.** Type III cell synapsing with 5 different nerve fibers. **D.** Type II cell forming synapses with 4 distinct nerve fibers. **E.** Example of a single nerve fiber (yellow; NF07) receiving synapses from different cell types. The Type II (blue; T2C06) cell forms channel synapses while the Type III cells (red; T3C06 & T3C07) form vesicular synapses onto the same fiber.

**Fig. 10.**
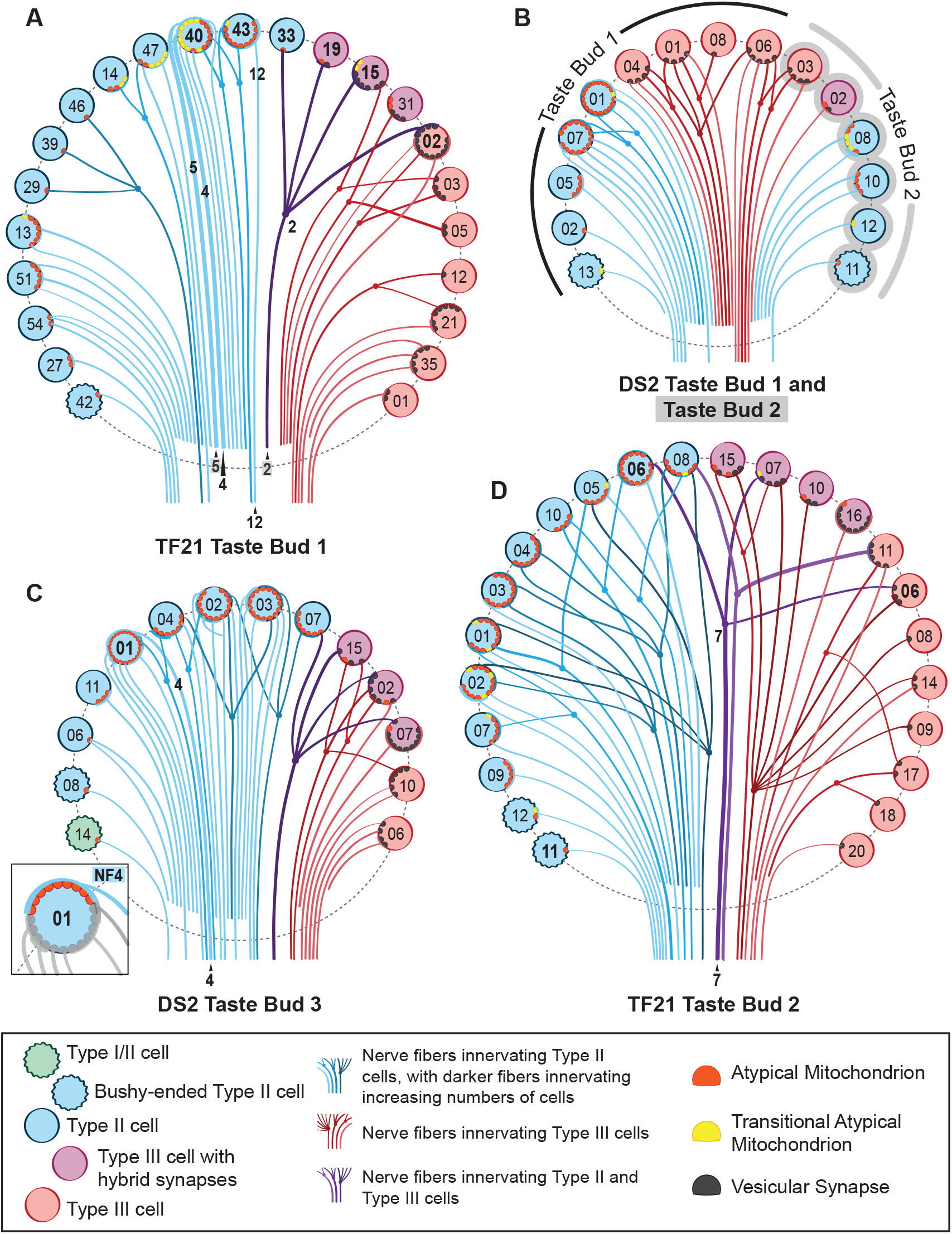
A connectome diagram for the taste buds characterized in this report. A) TF21 Taste Bud 1, B) DS2 Taste Bud 1 and Taste Bud 2 (conjoined), C) DS2 Taste Bud 3, and D) TF21 Taste Bud 2. Taste cells are denoted by colored circles as follows: green, wavy circle = Type I/II cell; teal, wavy circles = bushy-ended Type II cells; blue circles = Type II cells; purple circles = Type III cells with hybrid synapses; red circles = Type III cells with vesicular synapses. Each line denotes a single nerve fiber color coded: red fibers innervate Type III cells, blue fibers innervate Type II cells, Purple fibers innervate both Type II and Type III cells. Branched fibers are shown by dots and forked connections. Nerve fiber lines deepen in color for nerves that innervate increasing numbers of taste cells. Nerve processes that could not be traced to exit the taste bud are illustrated as not extending below the dashed line outlining each taste bud. Atypical mitochondria are denoted by red semicircles, while “transitional” atypical mitochondria are denoted by yellow semicircles. Vesicular synapses are denoted by black semicircles. Those cells mentioned in the text are denoted by bolded lettering, and nerve fibers mentioned in the text are labeled. The inset in C emphasizes a single nerve fiber that contacts 9 atypical mitochondria in T2C01—other features of this cell are greyed out to emphasize NF4.

While a single nerve fiber generally forms either all channel synapses with Type II cells or vesicular synapses with Type III cells, some uncommon nerve fibers form both types of connections with different cell types (Fig. 9E). Across all 5 taste buds in the sampling volume, of 88 total fibers forming channel synapses with Type II cells, only 4 fibers of these also form vesicular synapses with Type III cells. These 4 mixed synapse fibers also are a minority of the 44 total fibers receiving synapses from Type III cells. The probability of a taste bud having a mixed synapse fiber is directly related to the total number of Type II and Type III cells within the bud (Pearson’s correlation: r(3)= .89, p = .04). In several cases, a nerve fiber can form a channel synapse with a Type II cells and a vesicular synapse with a Type III cell, e.g. TF21_TB2 Nerve Fiber 7 and cells T3C06 and T2C06. Accordingly, the type of contact formed cannot be dictated entirely by the post-synaptic nerve fiber since a given fiber may participate in both types of synapses but on different cells. Furthermore, the convergence of Type II cell inputs and Type III cell inputs onto single nerve fibers shows that connectivity within a taste bud is not organized entirely on a labeled-line basis since Type II and Type III cells transduce different taste qualities.

An example of such cross-modal connectivity is NF2 in TF21_TB1, which forms a channel type synapse with a Type II cell (T2C33), a vesicular synapse on one Type 3 cell (T3C19), a complex series of channel synapses and 4 vesicle clusters on another Type III cell (T3C15) and 4 vesicular synapses with Type III cell T3C02, which also synapses onto 3 other nerve fibers (Fig. 10). Similarly, another nerve fiber in taste bud TF21_TB2 (NF07) receives a channel synapse from a Type II cell while also receiving vesicular synapses from 2 Type III cells (Fig. 9E). This shows that a given nerve fiber can form different types of synapses with different taste cells, and even different types of synapses with a single taste cell. A similar pattern of complex connectivity occurs for one fiber in taste bud DS2_TB3.

As can be appreciated from the connectome maps in Fig. 10, multiple branches and complex patterns of connectivity are not limited to fibers innervating different types of taste cells. For example, a single Type II cell (T2C40 in TF21_TB1), has 10 sites with channel synapses and these 10 sites form contacts with 8 different nerve fibers. Of the 8 nerve fibers, 6 are uniquely associated with this taste cell while 2 form channel synapses with but one other Type II cell (T2C43), which itself forms multiple synapses onto only one other nerve fiber (NF12). Further, each nerve fiber may be post-synaptic to multiple channel synapses from a single taste cell, e.g. NF5 receives 4 synapses from cell T2C40 sharing one of these sites with nerve fiber NF4. At the other end of the spectrum of connectivity is Type II cell T2C11 in TF21_TB 2 which contains but a single channel synapse connecting to a unique nerve fiber (Fig. 10).

##### Summary

In general, nerve fibers that receive synapses from a Type III cell are post-synaptic only to other Type III cells and conversely, nerve fibers post-synaptic to Type II cells generally are post-synaptic only to other Type II cells (Fig. 8). Nerve fibers innervating both Type II and Type III cells represent a little less than 5% (4 of 87) of the fibers innervating Type II cells but 9.1% (4 of 44) of those innervating Type III cells. These results suggest that a labeled-line type of connectivity is not absolute and that some fibers will carry an admixture of different quality information from Type II (sweet-umami-bitter) and Type III (sour) cells. We note that the limited cross-connectivity of Type II and Type III cells represents a lower limit for the degree of cross-modality information since we cannot identify convergence of different taste specificities of Type II cells, sweet, umami or bitter, since we do not have morphological markers for cells with these different taste receptors.

## DISCUSSION

Reconstructions of nerve fibers and taste cells from sbfSEM images reveal details about the anatomical organization of circumvallate taste buds thereby providing a foundational anatomical model (Swanson and Bota, 2010) for rational construction of hypotheses about physiological interactions of cells and neural networks within the system. The different anatomical types of cells within a taste bud directly relate to their function and chemical responsiveness. Accordingly, a detailed anatomical understanding of cellular interrelationships and interactions with nerve fibers will enable appropriately constrained models of the taste periphery.

Our principal conclusions relate to two widely-held concepts regarding the functional organization of taste buds (Ohla et al., 2019): 1) that taste information is transmitted to the brain over “labeled-lines”, i.e. nerve fibers conveying information about a single taste quality, and 2) that the different types of taste cells within a taste bud directly communicate with one another to modulate the transduction and transmission of taste information (Roper, 2007; Dando and Roper, 2009; Chaudhari and Roper, 2010; Vandenbeuch et al., 2020). These two concepts are interrelated in that cross-modal taste information could be generated by cross-talk between different types of taste cells even if the wiring were strictly according to cell type.

### Labeled-lines

The “labeled-line” hypothesis posits that the taste signal is quality-specific, i.e. a single nerve fiber will carry information about only a single taste quality, e.g. sweet or sour. The contravening hypothesis, the “across-fiber pattern theory”, holds that individual fibers respond to multiple qualities. Two recent studies directly comparing activity of taste ganglion cells in mice come to opposite conclusions about the responsiveness of the peripheral taste axons (Barretto et al., 2015)(Wu et al., 2015). Since Type III cells and Type II cells respond to different taste qualities, the labeled-line theory would predict that these different types of cells should synapse on entirely different populations of nerve fibers. Indeed, two recent studies suggest that at least some degree of specificity exists in the wiring of taste cells to taste nerves (Lee et al., 2017; Stratford et al., 2017). Conversely a study following branching patterns of nerves in fungiform taste buds finds that 28% of fibers contact both Type II and Type III cells (Huang et al., 2020). We find that about 9% of the fibers synapsing with Type III cells also receive synapses from Type II cells indicating that wiring specificity in circumvallate taste buds is not entirely consistent with the labeled-line model – some nerve fibers likely respond to stimuli activating the different taste cell types: sour for Type III cells (Huang et al., 2006; Huang et al., 2008; Kataoka et al., 2008) and either sweet, umami or bitter for Type II cells. Thus while connectional specificity may exist, it is not absolute.

### Cell-cell Communication

The adjacency of different cell types within a taste bud suggests the possibility of cell-cell intercommunication. Indeed, several studies indicate that neurotransmitters – ATP, serotonin, GABA, acetylcholine – released by one type of taste cell can influence the activity in nearby taste cells of similar or different types (Roper, 2013).

Accordingly, we had anticipated finding substantial, specialized areas of contact between the Type II and Type III cells as well as between adjacent Type II cells or Type III cells. Such is not the case. Type I cells effectively surround and largely isolate the Type II and Type III cells from one another. We found no specializations – synaptic or otherwise – at the limited points of membrane contact between these cells. The interactions between the different cell types observed in intact taste buds thus are likely due to either paracrine effects of neurotransmitter leaking past the Type I cells, or due to transcellular activation via Type I cells. The role of ectoNTPDase on Type I cells in modulation of activity of sweet responsive taste cells is well characterized (Dando et al., 2012; Kataoka et al., 2012). Similar Type I-cell mediated modulation of other taste transducing cells may be via release of transmitters similar to the non-synaptic release of gliotransmitters in the CNS (Newman and Zahs, 1998; Pascual et al., 2005; Araque et al., 2014).

Although Type I cells may play an important role in modulation of taste signaling within a taste bud, they show no membrane specializations typical of synapses at any areas of contact with nerve fibers or other cells. Many Type I cells form intimate contacts with nerve fibers, especially in the nuclear region (Yang et al., 2020), but display no accumulations of vesicles, membrane thickenings, or atypical mitochondria at the zones of contact.

### Synaptic Efficacy

The number of synapses made by a taste cell onto a single nerve fiber varies greatly and may be a determining factor for effectiveness of information transfer. Many taste cells – both Type II and Type III – form but a single synapse onto a single nerve fiber (Fig. 10). In contrast, other taste cells form multiple synapses onto a single nerve fiber; the maximum observed in our data is 10. Whether the number and size of the synaptic contacts relates to synaptic efficacy is unclear. Different Type II taste cells release different amounts of ATP to similar levels of chemical stimulation (Murata et al., 2010). Cells having more channel synapses may release more ATP than cells with fewer synapses, thereby activating the postsynaptic nerve fiber more effectively. A taste cell forming 10 channel synapses onto a nerve fiber would likely exert a much greater influence on the activity of that nerve fiber than would a cell with but a single synapse.

### Divergence and convergence

Substantial variety exists in terms of the number of taste cells contacted by individual nerve fibers and the number of nerve fibers synapsed onto by a single taste cell. In all but the smallest taste buds, at least one nerve fiber in each bud branches within the taste bud to connect synaptically with 4 taste cells, often including both Type II and Type III cells. This suggests that these fibers are broadly responsive to both sour (Type III) and one or more GPCR-mediated taste qualities, i.e. bitter, sweet or umami (Type II). Such multiple responsiveness is reported for both gustatory nerve fibers and ganglion cells (Barretto et al., 2015; Wu et al., 2015). Further, since fibers likely branch before entering a taste bud, a fiber innervating Type II cells in one taste bud may innervate Type III cells in another bud, but we would not be able to detect such convergence with the methods employed. Furthermore, with the methods employed, we would be unable to detect convergence of Type II cells of differing specificities unto single nerve fibers. Hence our population counts yield only a lower limit on the degree of convergence likely in this system.

The number of axons postsynaptic to any given taste cell varies from 1 – 8. How this variability affects the output of the system is unclear. A multiply-connected taste cell might exert a larger influence on total nerve output than would a taste cell synaptic onto but a single fiber. If the presynaptic cells are of different taste specificity, then the more connected cell may drive the response of the nerve fiber at low concentrations of tastant whereas the activity of the less connected cell becomes evident only at higher concentrations. Indeed, differences in specificity of response with concentration (Wu et al., 2015) is reported for taste ganglion cells. A ganglion cell may respond to a single taste quality at low concentrations, but show broader responsiveness at high stimulus concentrations.

### Diversity of Type II Cell Morphology

Type II cells, those transducing sweet, umami or bitter tastes have been conventionally characterized as elongate, smooth cells having a large, ovoid nucleus with sparse heterochromatin and channel-type synapses with “atypical” mitochondria (Clapp et al., 2004; Kinnamon and Yang, 2008; Yang et al., 2020). In our previous work, we showed that, in mice, such cells terminate apically with a single, stout microvillus (Yang et al., 2020). In our current study, we identify a subset of bushy-top Type II cells -- about 14% of the population -- that end below the taste pore with an irregular assembly of short microvilli. While these may correspond to a unique functional class of taste cell, we suggest they may be representative of immature Type II cells. Type II cells have a life span of 8-20 days (Perea-Martinez et al., 2013) with markers of transduction, gustducin, PLCß2 or IP3R3 first appearing around days 2-3 (Cho et al., 1998; Miura et al., 2006; Koyanagi-Matsumura et al., 2021). Accordingly, about 10-12.5% of Type II cells in any taste bud will be in an immature state. The bushy-top Type II cells each form a channel synapse with but a single nerve fiber and have only a single specialized point of contact with that fiber. If these bushy-top Type II cells represent an immature morphological state, then this suggests a possible order of maturation where a synapse forms first followed by full elaboration of the apical end of the cell in the form of a single stout microvillus reaching into the taste pore.

## ACKNOWLEDGEMENTS

The authors thank Dr. Grahame Kidd and Emily Benson for acquisition of the sbfSEM datasets, and Dr. Sue Kinnamon (Univ. Colorado Sch. Medicine) for comments on drafts of this work.

